# Environmental Impacts of Dry Vegan Dog Food: A Comprehensive UK Analysis

**DOI:** 10.1101/2025.05.19.654875

**Authors:** R.A. Brociek, D.S. Gardner

## Abstract

**Introduction:** Pet food production contributes substantially to global environmental pressures, driven largely by animal-derived ingredients. The current study quantified the environmental impacts of 31 commercially available dry dog foods purchased in the UK, categorised as plant-based, poultry-based, red meat-based (beef and lamb) and veterinary renal diets.

**Method:** Environmental metrics including land use (m²/1000 kcal), greenhouse gas emissions (kg CO₂eq/1000 kcal), acidifying emissions (g SO₂eq/1000 kcal), eutrophying emissions (g PO₄³⁻eq/1000 kcal), and freshwater withdrawal (L/1000 kcal) were estimated using life cycle assessment datasets and adjusted for ingredient composition and energy density. Foods were additionally standardised to 100% moisture-adjusted content, and group differences were evaluated statistically.

**Results and discussion:** Plant-based diets demonstrated the lowest impacts across all measures. Poultry-based and veterinary diets showed intermediate profiles, while beef- and lamb-based foods exhibited substantially higher impacts. Beef-based diets required 101.12 m² land and emitted 31.16 kg CO₂eq per 1000 kcal dry food, compared to 2.71 m² land and 2.79 kg CO₂eq for plant-based. Beef-based foods generated 7.1-fold higher acidifying emissions and 16.4-fold higher eutrophying emissions, compared to plant-based foods. Lamb-based foods were the most freshwater intensive (677.01L/1000kcal), compared to plant-based (246.95L/1000kcal). Whilst moving to a circular food system has the potential to reduce environmental impact, current pet food manufacturing practices compete with human food supply and are not nutritionally necessary for dogs to thrive. These findings highlight a major opportunity to mitigate the environmental footprint of companion animal nutrition through reformulation towards higher inclusion of plant-based ingredients.

## Introduction

As consumers seek more sustainable alternatives to eating traditional, meat-based products, the popularity of going ‘plant-based’ has increased, with many owners wishing to feed their pets a more sustainable pet food (Rothgerber, 2013; Ingenpaß *et al*., 2021). Expansion of the population of humans on the planet has inevitably placed demands on the food chain, despite increased production efficiencies, resulting in a number of environmental effects; increased greenhouse gas emissions, deforestation, land degradation, biodiversity loss, freshwater depletion and pollution of soil and water sources (Pimentel and Pimentel, 2003; Reijnders and Soret, 2003; Westhoek *et al*., 2014; Poore and Nemecek, 2018; Humpenöder *et al*., 2022; Scarborough *et al*., 2023).

In 2018, the global canine population was estimated at 471 million (Joe Sivewright and Nina Leigh Krueger, 2019). Feeding these omnivores a meat-based diet, even when accounting for the use of ‘by-products’, generates considerable greenhouse gases (GHG). The potential for exchanging meat-for plant-based alternatives, when feeding companion animals, could therefore, significantly reduce the environmental impact of pet food (Humpenöder *et al*., 2022). A number of previous studies have assessed the environmental impact of pet foods (Swanson *et al*., 2013; Okin, 2017; Su and Martens, 2018a; Su, Martens and Enders-Slegers, 2018; Martens, Su and Deblomme, 2019; Pedrinelli *et al*., 2022; Jarosch, Bach and Finkbeiner, 2024). The estimated environmental impact of manufacturing dog food in Brazil has been suggested to account for 2.9 – 24.6 % of the total carbon dioxide emissions of the country (Pedrinelli *et al*., 2022). In China, it was estimated that dogs fed commercially-produced pet food accounted for three-fold greater carbon emissions than dogs solely eating leftover, human-grade food (Su, Martens and Enders-Slegers, 2018). In 2015, the GHG footprint of cats and dogs in China – in terms of food consumption – was estimated to be between 2.5 – 7.8% of the GHG footprint of the total population in China, which is not inconsiderable. Globally, estimates suggest that production of pet food, including use of by-products, requires 41-58 million hectares (mHa) of agricultural land, around twice the land area of the UK (24.9mHa), producing equivalent GHG emissions of whole countries such as Mozambique or the Philippines (Alexander *et al*., 2020).

Producing animal protein (e.g. beef and lamb) for human (or pet) consumption, registers, without doubt, the greatest impact on the environment in terms of measures including land-area, water-use and GHG emissions. Producing plant-based protein records a fraction of the environmental cost; indeed, even when comparing pea-protein producers with the highest carbon footprint (i.e. 0.8lkg CO_2_eq/100lg protein) versus the lowest impact farm for lamb, beef or chicken production (12.0, 9.0 and 2.4lkg CO_2_eq/100lg protein, respectively), the difference remains particularly stark, particularly if this is scaled-up globally (Poore and Nemecek, 2018; Ritchie, 2020). Although studies have investigated the reduced environmental impact of people adopting a plant-based dietary pattern, and the reduced impact of plant-based pet food consumption by companion animals, none have directly compared the environmental impact of meat versus plant-based (vegetarian and vegan) dog food in the UK. Jarosch, Bach and Finkbeiner (2024) estimated the environmental impact of wet, vegan dog food. However, wet foods are known to have a higher environmental impact than dry foods (Pedrinelli *et al*., 2022; Jarosch, Bach and Finkbeiner, 2024) and 50-70% of UK pet owners choose to feed dry kibble (Morelli, Stefanutti and Ricci, 2021; Pet Food Industry, 2023), either on its own or mixed with wet. Previous studies evaluating the impact of pet food manufacturing in Japan (Su and Martens, 2018b) and China (Su, Martens and Enders-Slegers, 2018) are not directly applicable to the UK given our variations in pet preferences and population, as well as access to different ingredients. Additionally, other studies (Swanson *et al*., 2013; Okin, 2017; Alexander *et al*., 2020; Knight, 2023) have calculated the environmental impact of dog foods *per se* using a variety of different methods.

In the current study, using validated methods to assess and estimate the environmental footprint of pet food production, we estimate the extent of the reduced impact of feeding plant-based versus other common diets fed in the UK (meat – beef, lamb; veterinary-diets – semi-synthetic animal-based ingredients).

## Methods

The environmental impact of 31 adult dog foods commercially available in the UK, were calculated with regards to land use (m^2^/1000kcal), carbon dioxide equivalent (CO_2_eq(kg)/1000kcal), acidifying emissions (SO_2_eq(g)/1000kcal), eutrophying emissions (PO_4_ ^3-^eq(g)/1000kcal) and freshwater withdrawal (L/1000kcal). These were calculated using previously published values (Poore and Nemecek, 2018).

### Selection of dog food

Thirty-one complete dry dog food samples were acquired from VioVet online, Pets at Home and supermarkets in the UK, representing twenty-seven different brands. Exclusion criteria included not being a ‘complete’ food, not being labelled as for either ‘adult’ or ‘adult and senior’ dogs, foods that weren’t readily available to the public (i.e. required a prescription) and foods that did not come in packages of 3kg or less, to limit food waste. Foods were grouped according to their main protein source, “meat-based”; poultry (n = 7), lamb (n = 6) and beef (n = 6) as the main listed ingredient or “plant-based”; which included vegan (n = 4) and vegetarian foods (n = 2). In addition, n=6 veterinary diets were tested. These “veterinary” diets were all specifically formulated for renal patients. These had no specific flavour claimed and no named meat ingredient but had a guaranteed analysis on the label. Since they are largely comprised ‘synthetic’ and thus highly controlled ingredients, rather than being “meat- or plant-based” then they were treated as a discrete group. Pork, rabbit and other meats were not included in the current study as the market for these products is limited in the UK, with few products available with these as the sole meat source. Fish was also not included due to calculations, particularly land use, being incomparable to terrestrial animals. Foods for the testing were procured over a 4-week period.

### Pet food composition and ingredient weightings

Across 31 foods, all 298 ingredients were reviewed, encompassing up to 33 individual ingredients. Most individual foods (27/31, 87%) included between 6 and 20 ingredients. All ingredient terms used were pre-defined (Pedrinelli *et al*., 2022). In summary, “meats” are considered “raw meat” unless specified on packaging and “meat meals” are considered to be cooked. “Animal protein hydrolysate” is reported as “chicken protein hydrolysate”. Vegetable percentages are provided by fresh weight where possible. “Animal fat” (where the animal is not specified) and “poultry gravy” have been reported as “poultry fat”. “Dehydrated poultry protein” has been reported as “poultry by-product meal”. Where “oils & fats” are listed, this is reported as “sunflower oil, unspecified”. “Salmon oil” and “fish oil” are both reported as “fish oil”. Rye and spelt are reported as “whole wheat”. Maize gluten has been reported as “corn gluten meal”. “Wheat” has been reported as “wheat gluten”. All varieties of wheat have the same impact values. Unless listed in the ingredients as “brown rice”, rice has been reported as “white rice, cooked”. Pasta is reported as “spaghetti”. All “potato protein” is reported as “potato starch”. Potato starch and cooked potato have the same impact values. Where label ingredients did not have an equivalent ingredient with pre-calculated impact value, these were excluded from the estimation. Excluded ingredients include, but were not limited to, glycerol, minerals, lupine, alfalfa, algae, botanicals, vegetable stock and chicory. In 30/31 foods, the first 4 ingredients, which is often most of the food composition, were always included in the estimation. This is with the exception of one food with “glycerol” as the third ingredient. Where “animal by-product” (ABP) is referred to, this includes but is not limited to hides, skins, hydrolysed protein, rendered fat, digest, milk by-products, eggs, fish products and bones (Gov.uk, 2019).

Previously, ingredient weightings were based on the first five ingredients, each given a 20% weighting, or a 1.5 ratio in descending order of ingredient inclusion (e.g., 38.4% for the first ingredient, 25.6% for the second, 17.1% for the third, etc… (Okin, 2017; Alexander *et al*., 2020). In the present study, a modified ratio, based on the dietary group being analysed, provided a more accurate representation: ingredient percentages were taken from package ingredient lists, assuming a descending order of percentage incorporation (see Supplementary Information). When proportions were undefined, an average percentage for the dietary group (e.g., meat-based, plant-based, or veterinary) was applied based on the ingredient’s position in the list (e.g., fifth ingredient). In cases where ingredients were listed as a single percentage (e.g., “dried chicken and turkey 30%”), a 50:50 ratio was used (15% dried chicken, 15% dried turkey). Using this method, the total ingredient composition ranged from 39.29% to 120%, before being adjusted to 100% to ensure an equitable comparison between all foods.

### Environmental impact assessment

Life cycle assessments (LCA) are the current standard for assessing environmental impact, with many studies using some variation (Van Zanten *et al*., 2016; Yavor, Lehmann and Finkbeiner, 2020; Jarosch, Bach and Finkbeiner, 2024). Estimations of five environmental impact scores (per 1000kcal metabolisable energy [ME]) were calculated for all pet foods, including 1) land use (m^2^) as shown below in Eq. 1, and 2) carbon dioxide equivalent (CO_2_eq), 3) acidifying sulphur dioxide (SO_2_), 4) eutrophying phosphate (PO_4_ ^3-^) emissions and 5) freshwater withdrawal (L), with the only difference for additional calculations being the appropriate impact value (Pedrinelli *et al*., 2022) in place of “ingredient land use impact value” (Eq. 1).

Impact values represent a methodologically harmonised dataset of life cycle assessments (LCA) from 570 suitable studies: representing ∼38,700 farms in 199 countries and producing ∼90% of the global protein and calories (Pedrinelli *et al*., 2022). Over 1000 post-farm processes were included in the production of these impact values, including processing, packaging, retail, transport and losses, up the point of consumer choice.

Moisture content of the food from each food was used to determine its final energy density. Final impact values (/1000kcal ME) were determined considering ingredient percentage composition, the impact value of each ingredient and the food moisture content. Finally, a factor was applied to standardise all foods to a theoretical 100% inclusion.

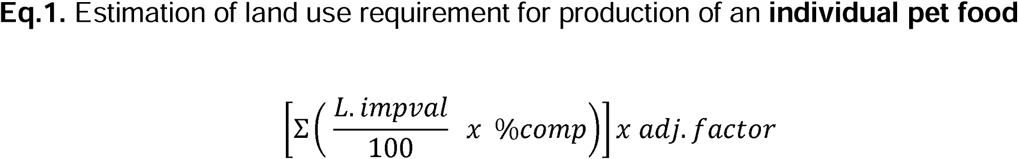

where L.impval (land impact value), expressed in m^2^/1000kcal, was previously calculated (Pedrinelli *et al*., 2022), %comp is the moisture-adjusted percentage composition of ingredient in the food. For each food, the sum impact of all individual ingredients was totalled, and an adj.factor was the value applied to adjust the total food ingredient composition to 100%, calculated based on the listed and theoretical percentage composition of ingredients. To calculate carbon dioxide equivalent (kg CO_2_eq/1000kcal), acidifying emissions (g/1000kcal SO_2_eq), eutrophying emissions (g/1000kcal PO_4_ ^3-^eq) and freshwater withdrawal (L/1000kcal), respective impact values were substituted. These were replaced in the equation as appropriate to determine the impact of each parameter, for each ingredient and subsequently, each food.

### Statistical analysis

Data were analysed using analysis of variance (ANOVA) for the fixed effect of the three diet-types (meat-based or plant-based, veterinary). To meet assumptions for analysis by ANOVA, all data were checked for a normal distribution of residuals and respective Q-Q plots. If necessary, non-normally distributed data were log-transformed (log_10_) prior to analysis by ANOVA or a suitable non-parametric, distribution independent test was used (e.g. Kruskall-Wallis NP-ANOVA). All such data were analysed using GraphPad Prism v10.3.0 (GraphPad Software Inc., California, USA) and GenStat v22 (VSNi Ltd., Rothamsted, UK).

### Data availability

All anonymised data for the products used in this manuscript are available from The University of Nottingham research data repository.

## Results

Land use (m^2^/1000kcal), carbon dioxide equivalent (kg CO_2_eq/1000kcal), acidifying emissions (g/1000kcal SO_2_eq), eutrophying emissions (g/1000kcal PO_4_ ^3-^eq) and freshwater withdrawal (L/1000kcal) were calculated for each ingredient of every food, individually (Table 1). These values were adjusted for the relative percentage of the food that the ingredient takes up, using listed and theoretical proportions (see Materials and Methods for further details). Data for each food were then calculated per 1000kcal fed to account for differences in moisture content and energy density between foods.

**Table 1.**
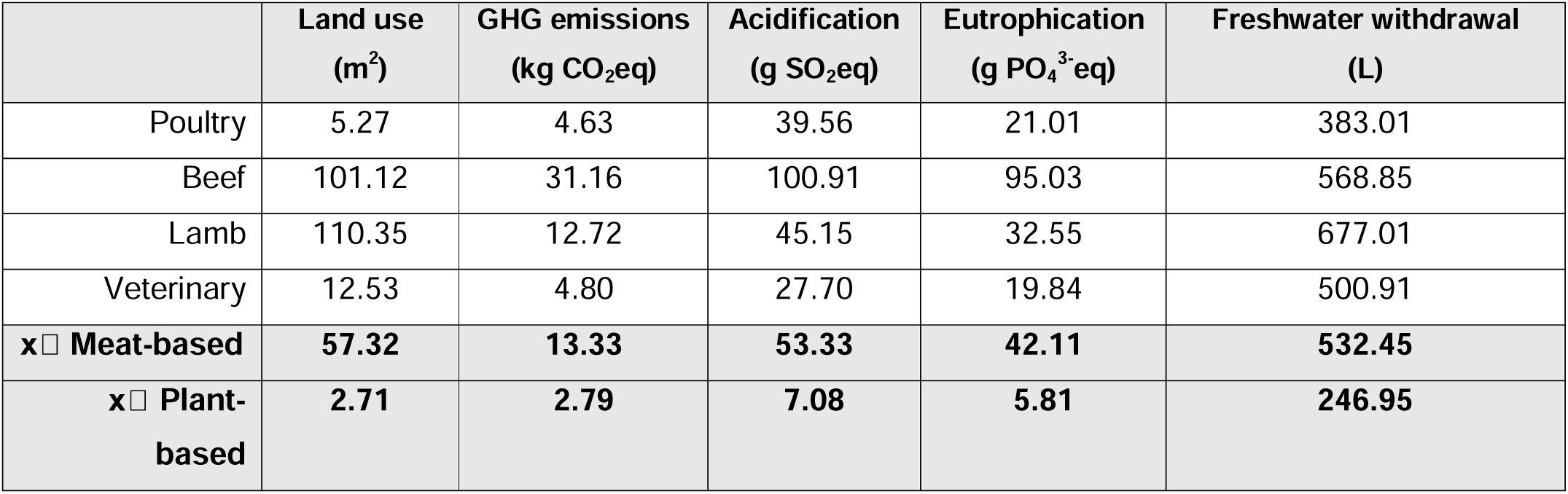
Estimated average land use, GHG emissions, eutrophication, terrestrial acidification and freshwater withdrawal of dry kibble dog foods, per 1000kcal as fed.

|  | Land use<br>(m<sup>2</sup>) | GHG emissions<br>(kg CO<sub>2</sub>eq) | Acidification<br>(g SO<sub>2</sub>eq) | Eutrophication<br>(g PO<sub>4</sub><sup>3-</sup>eq) | Freshwater withdrawal<br>(L) |
| --- | --- | --- | --- | --- | --- |
| Poultry | 5.27 | 4.63 | 39.56 | 21.01 | 383.01 |
| Beef | 101.12 | 31.16 | 100.91 | 95.03 | 568.85 |
| Lamb | 110.35 | 12.72 | 45.15 | 32.55 | 677.01 |
| Veterinary | 12.53 | 4.80 | 27.70 | 19.84 | 500.91 |
| **x □ Meat-based** | **57.32** | **13.33** | **53.33** | **42.11** | **532.45** |
| **x □ Plant-based** | **2.71** | **2.79** | **7.08** | **5.81** | **246.95** |

Overall, plant-based foods always had the lowest environmental impact scores for each parameter, and beef and/or lamb always had the highest (Figure 1a-e). Veterinary foods, being semi-synthetic, consistently scored between these two extremes of foodstuff. When combined in an unbiased, multivariate analysis lamb and beef-based foods were clearly discriminated along the first and second principal components from all other foodstuffs, which were broadly similar. These corresponded, with the highest discriminant scores, to the combination of land-use, freshwater withdrawal and CO_2_eq production for lamb (contributing significantly to PC1) and to CO_2_eq + SO_2_eq + PO4^3-^eq emissions for beef (contributing significantly to PC2; Figure 1f). Poultry had significantly lower environmental impact than beef or lamb and co-located with plant-based foods in multivariate analyses of centred-data for all impact measures (Figure 1f).

**Figure 1.**
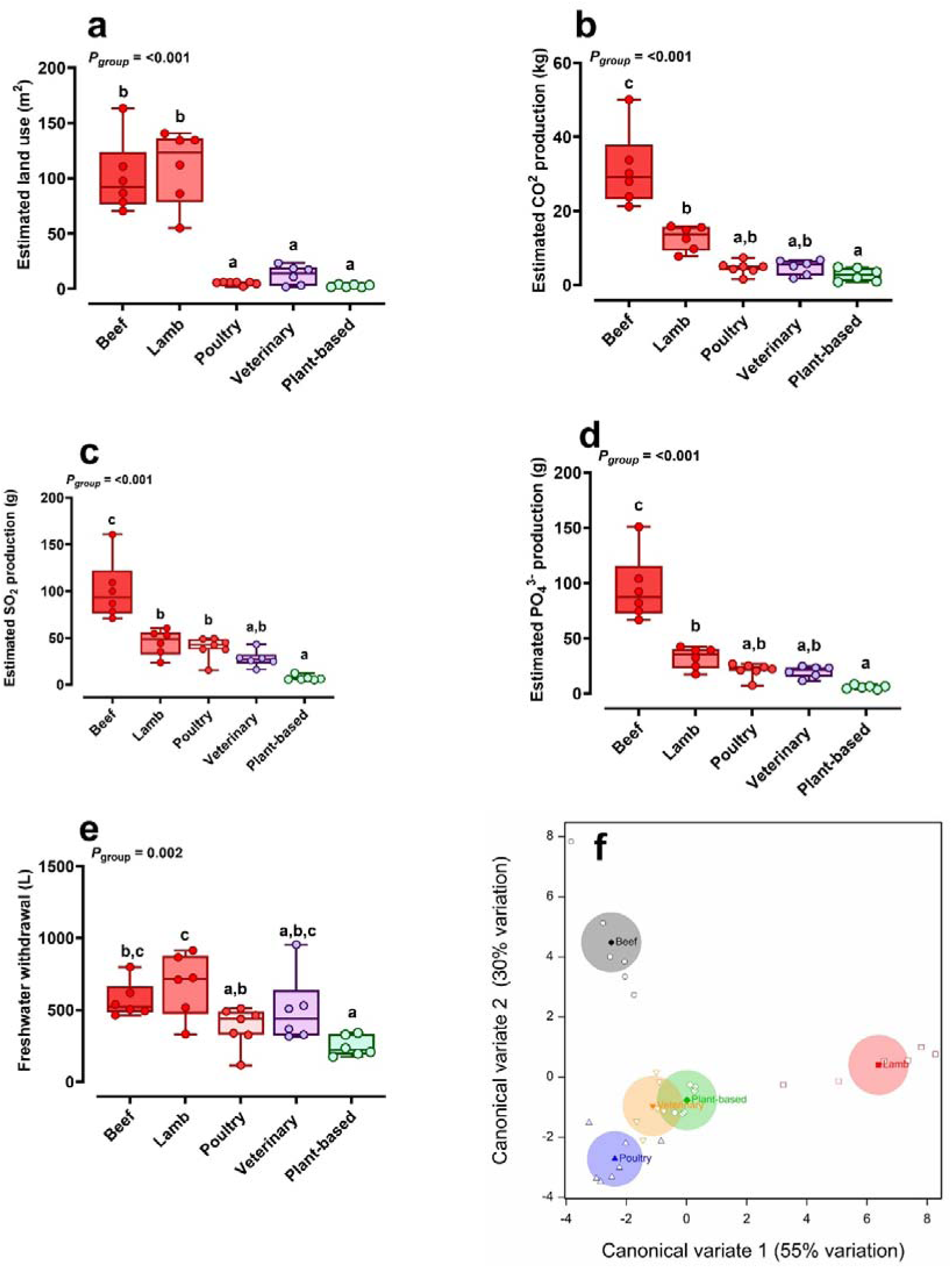
Estimated environmental impact of dry feeds for dogs according to feed-type. **a-e,** individual data points and box-plots (line at median, box = first to third interquartile range, whiskers = min, max range) for estimates of environmental impact. Equations used to derive values for each feed as described in Methods (Eq 1; land-use) or supplementary information (Eq 2-5). Type of meat used to produce ‘meat-based’ food presented for clarity. Veterinary diets are mostly synthetic (see Methods) **f),** multivariate analysis (canonical discriminant plot) with all feeds (n=31) and all estimated environmental impact data (as standardised z-scores due to the marked differences in numbers) represented.

### Food composition

From the labels, the average content of proximates (protein, fat and fibre) of meat- and plant-based foods were similar (meat: 23.50%, 12.78% and 2.9% versus plant: 24.63%,10.45% and 4.00%, respectively). Veterinary foods had lower protein (by design) with higher fat inclusion (13.82%,17.28% and 2.4%, respectively).

Across all foods, 60 different ingredients (after removal of ingredients without suitable impact values to apply) were assessed across the 31 foods. Almost one third of ingredients were animal-derived (19/60, 32%). When comparing the top 5 ingredients of all foods, the most common ingredients were corn gluten (present in 19/31 foods), white rice (12/31), poultry fat (10/31), sunflower oil (7/31), wheat gluten (6/31) and brown rice, raw chicken meat, meat and bone meal, pea, potato and textured vegetable protein (all 5/31). When taking the full formulation into account (up to 21 ingredients), the top ingredients were very similar; corn gluten meal (22/31), poultry fat (17/31), white rice (14/31), sunflower oil (15/31), flaxseed and brewer’s yeast (both 12/31) and pea (10/31).

Across the 25 meat-based foods there were 54 different ingredients, 19 of which were animal-derived. The four most common ingredients were poultry fat (18/25), corn gluten meal (17/25), beet pulp and white rice (both 12/25). The 6 plant-based foods contained 26 different ingredients, with one having an animal-derived ingredient – egg powder. The four most common ingredients were sunflower oil and brewer’s yeast (both 6/6), corn gluten meal and pea (both 5/6).

### Land use

The average land use (Figure 1a) required to produce lamb and beef dog foods were similar (*P* = 0.70), as were poultry and veterinary foods (*P* = 0.20). However, the difference between the pairs and between these groups versus plant-based were statistically significant (*P* = <0.001 for plant-based vs beef, lamb and veterinary, and *P* = 0.01 plant-based vs poultry). The average estimated land use required to manufacture each dry dog food was 110.35m^2^/1000kcal (range: 54.9 – 140.4) for lamb, 101.12m^2^/1000kcal (range: 70.3 – 163.0) for beef, 5.27m^2^/1000kcal (range: 2.13 – 6.26) for poultry and 2.71m^2^/1000kcal (range: 1.51 – 4.02) for plant-based. Veterinary diets required 12.53m^2^/1000kcal (range: 1.88 – 23.36), aligning with poultry and plant-based foods, but remaining statistically significantly different (*P* = 0.04 and 0.01, respectively). Veterinary foods differed significantly from beef (*P* = <0.001) and lamb (*P* = <0.001). Individual ingredients recording greatest land use included lamb meal (92.88m^2^/1000kcal), meat and bone meal of unknown species (47.18m^2^/1000kcal) and beef liver (25.17m^2^/1000kcal) (Figure 2). Individual ingredients recording greatest land use in plant-based foods were primarily soy, such as soy protein concentrate (1.67m^2^/1000kcal) and cooked soy (1.87m^2^/1000kcal).

**Figure 2.**
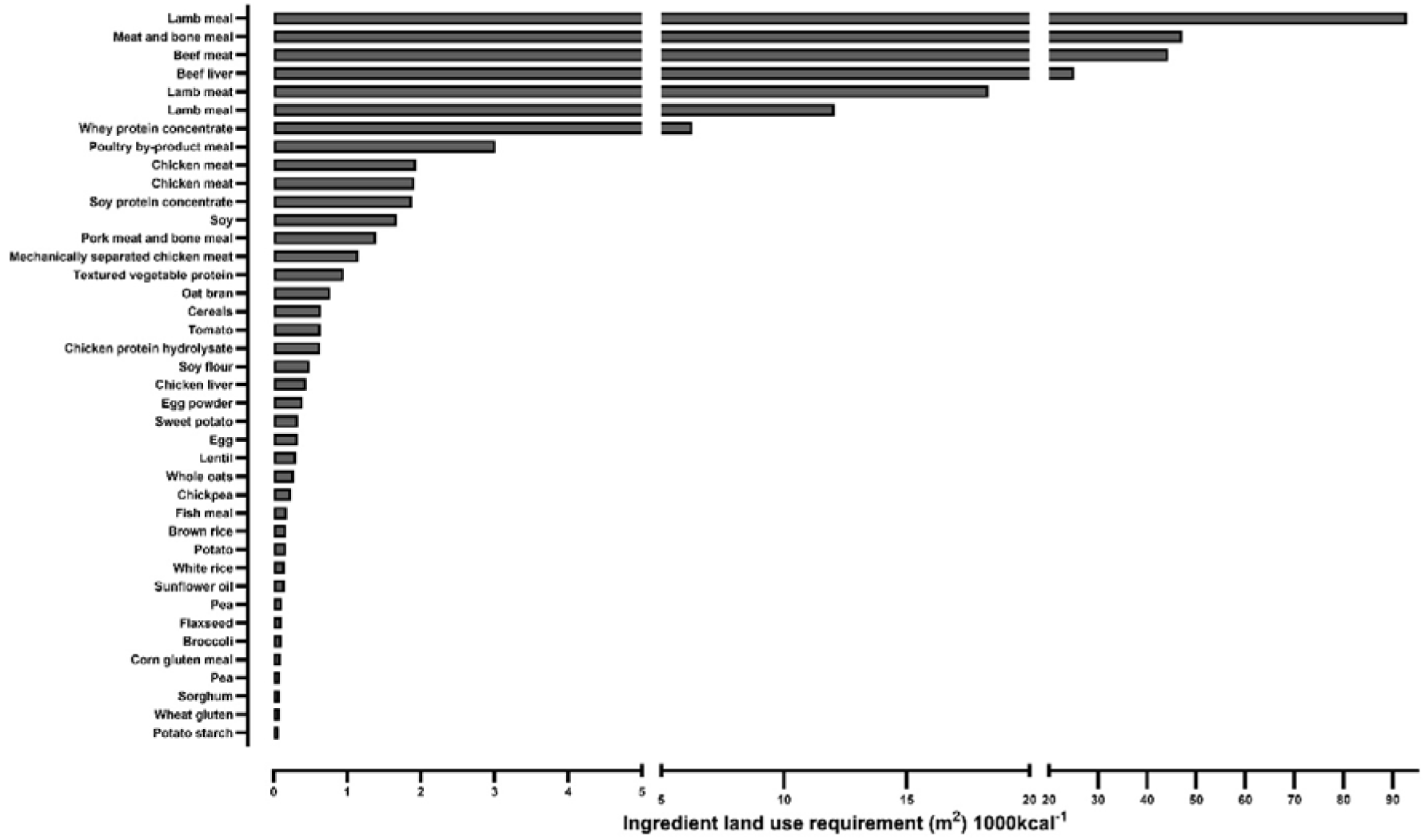
Average land use requirement (m^2^ 1000kcal^-1^ of dry food) of top 40 ingredients used in the current study.

### Carbon dioxide equivalent (CO_2_eq) emissions

In agricultural production, greenhouse gases include carbon dioxide, methane and nitrous oxide. To comprehensively capture all GHG emissions from dog food production, emissions were expressed as carbon dioxide equivalents (CO_2_eq) in kg CO_2_eq/1000kcal of food. This metric accounts for the global warming potential (GWP) of each gas. GWP100 is a measure of the relative warming impact one molecule or unit mass of a greenhouse gas, relative to carbon dioxide (which has a GWP of 1), over 100 years (Allwood J.M., V. Bosetti, N.K. Dubash, L. Gómez-Echeverri, and C. von Stechow, 2014; Ritchie, Rosado and Roser, 2022). CO_2_eq values reflect emissions across the entire supply chain, including fertiliser quantity and type, irrigation use, soil, and climatic conditions.

Plant-based foods had the lowest GHG emissions (2.79kg CO_2_eq/1000kcal) while poultry (4.63kg CO_2_eq/1000kcal) and veterinary foods (4.80kg CO_2_eq/1000kcal) were non-significantly different (*P* = 0.09 and *P* = 0.12, respectively) (Table 1). Beef (31.16kg CO_2_eq/1000kcal) was significantly higher than both plant-based (*P* = <0.0001) and poultry (*P* = <0.0001) diets. On average, lamb foods (12.72kg CO_2_eq/1000kcal) were lower than beef, and more similar to poultry and plant-based foods, than they were to beef.

Ingredients with highest CO_2_eq impact values for meat-based foods included meat and bone meal (99.78 CO_2_eq/1000kcal), raw beef liver (99.69 CO_2_eq/1000kcal), beef liver (49.120 CO_2_eq/1000kcal), lamb meal (37.35 CO_2_eq/1000kcal) and whey protein concentrate (28.16 CO_2_eq/1000kcal). The highest impact values in plant-based ingredients were pea, unspecified (13.31 CO_2_eq/1000kcal), pea, cooked (12.72 CO_2_eq/1000kcal), tomato, raw (5.62 CO_2_eq/1000kcal) and wheat gluten (1.40 CO_2_eq/1000kcal).

### Eutrophying and Acidifying emissions

Eutrophying emissions (PO_4_^3-^eq) is a measure of the pollution of soil and waterways from nutrient-rich runoff, particularly inorganic phosphorus (used as a fertiliser) and nitrogen dioxide (Ritchie, Rosado and Roser, 2022). Acidifying emissions (SO_2_eq) groups sulphur dioxide (SO_2_), nitrogen oxides (NOx) and nitrates from fertilisers and other stages of the manufacturing process (waste water, animal feedlots etc.), which react with water and oxygen to form acidifying compounds such as sulphuric and nitric acids in the atmosphere, before falling as acid rain (Reijnders and Soret, 2003; US Environmental Protection Agency, 2024).

For both metrics (Table 1), plant-based food produced the lowest emissions, and beef the most (Figure 1c,d). Eutrophying (PO_4_ ^3-^eq) values were significantly different when comparing plant-based vs poultry (*P* = 0.0001) and plant vs veterinary, lamb and beef (*P* = <0.0001). Similarly, plant-based acidifying (SO_2_eq) values were significantly lower than veterinary (*P* = <0.001) and all meat-based foods (*P* = <0.0001). When comparing meat-based foods, beef foods scored significantly higher for both SO_2_eq and PO_4_ ^3-^eq (beef compared to poultry: [SO_2_eq] *P* = <0.001 and [PO_4_ ^3-^eq] *P* = <0.0001, lamb: [SO_2_eq] *P* = <0.01 and [PO_4_ ^3-^eq] *P* = <0.001, veterinary: [SO_2_eq] *P* = <0.001 and [PO_4_ ^3-^eq] *P* = <0.0001). Acidifying beef values were 2.2, 2.6, 3.6 and 7.1 times that of lamb, poultry, veterinary and plant-based foods, respectively, and eutrophying emissions were 2.9, 4.5, 4.8 and 16.4 times higher.

### Freshwater withdrawal

Total freshwater calculations refer to the freshwater used in agriculture and industry during the life cycle of pet food manufacture, including water used for irrigation, livestock production and for direct industrial use (including withdrawals for cooling thermoelectric plants) (World Bank Group, 2025). Lamb-based diets had the highest freshwater withdrawal (677.01L/1000kcal). Beef was lower than lamb, but nevertheless high (568.85L/1000kcal) as were veterinary foods (500.91L/1000kcal) (Figure 1e), relative to plant-based foods and poultry. The latter had the second lowest freshwater use (383.01L/1000kcal) with plant-based diets the least (246.95L/1000kcal). Hence, plant-based food had significantly lower freshwater withdrawal than lamb (*P* = <0.01), beef (*P* = <0.0001) and veterinary foods (*P* = <0.01).

## Discussion

This study is the first to comprehensively compare the environmental impact of commercially available meat-based and plant-based dry dog foods available on the UK market. Our findings broadly align with global literature, confirming that plant-based diets are consistently the lowest-impact choice across metrics of land use, greenhouse gas emissions, eutrophication, acidification, and water use (Pedrinelli *et al*., 2022; Knight, 2023). In contrast, diets containing lamb or beef generated the highest environmental burden, followed by veterinary and poultry-based foods. These differences have substantial implications when scaled to the lifetime of a single dog, as well as across the broader UK pet population, or indeed the entire planet.

A dog is considered adult for approximately 7–11 years (dependent on breed), preceded by the growth and development (puppy) stage and followed by the senior stage, with some also going through pregnancy and lactation. Foods specific for each life stage provide dogs with nutrition appropriate to their body’s needs at that point in life. In this study, the authors only evaluated adult dog food, and so estimations can only be extrapolated for this life stage. The environmental impact of diet varies depending on ingredient composition, with substantial differences between meat-based and plant-based formulations.

A 20 kg Labrador Retriever, the most popular dog in the UK currently, with low to moderate exercise, requires approximately 280 g of food (1008 kcal) per day. Over nine years, this amounts to approximately 919.8 kg of food (280 g/day × 365 days × 9 years). Conventional diets show a fold-increase in impacts across key environmental metrics when compared to plant-based alternatives (Figure 3). Over the 9 adult years, exclusively feeding a 20 kg Labrador Retriever plant-based, veterinary, poultry, beef, or lamb food would require 8,964 m² (about 1.4 football fields; each field is approximately 100 m x 64 m), 17,453 m² (around 2.7 football fields), 41,476 m² (approximately 6.5 football fields), 334,851 m² (roughly 52 football fields), and 365,409 m² (close to 57 football fields) of land, respectively. These results are largely supported by those currently in the literature (Robert Vale and Brenda Vale, 2009; Martens, Su and Deblomme, 2019).

**Figure 3.**
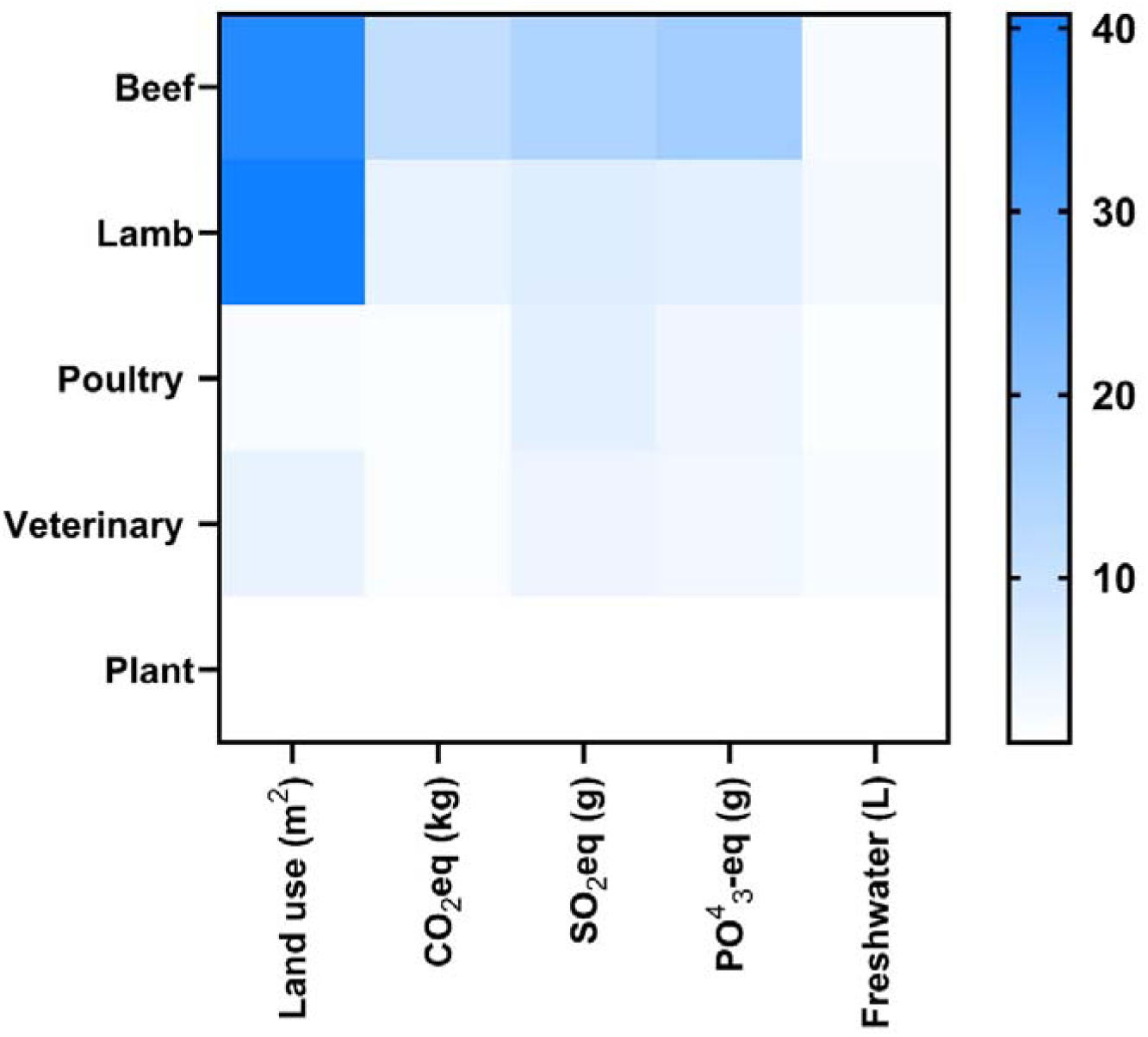
Heat map of fold-increase, estimated impact value for all categories of foods fed over a nine-year period (average length of time an adult diet is fed), compared to feeding plant-based.

Correspondingly, the greenhouse gas emissions (kg CO₂eq) produced by the plant-based, veterinary, poultry, beef, or lamb foods over this period would be equivalent to 2.8, 4.65, 4.8, 12.8, and 31.3 round trips between London and New York, per passenger, on a Boeing 747. These values are similar to results previously described (Yavor, Lehmann and Finkbeiner, 2020). Impacts also vary significantly depending on the size of the dog, and its energy needs (Satriajaya, 2017). Vale and Vale (2009) estimated that feeding a 15kg dog 300g/day of meat-based food requires 0.27 ha/year, whereas a Chihuahua would require 0.09 ha/year, a Scottish Terrier 0.18 ha/year, and a large Alsatian 0.36 ha/year. Scaling this across millions of dogs adds considerable pressure to land systems already under stress from both human and livestock food production.

### Land Use

Land use emerged as one of the most contrasting indicators of environmental impact among different dog food formulations. Our study’s land use estimates incorporated several refinements to increase accuracy, including multicropping (up to four harvests per year), periods of fallow (uncultivated land between crops), and economic allocation of co-products such as straw (Poore and Nemecek, 2018). These adjustments reflect real-world farming systems more accurately than older models, but they still highlight the large disparity in land demand between animal-based and plant-based dog food.

In this study, plant-based diets required an average of just 2.71 m² per 1000 kcal, compared with over 100 m² for beef and lamb-based diets. With soy and soybean concentrate having the highest plant-based impacts, the current study does support claims of soy products having a high land impact, with the soy crop production utilising over 280 million acres used globally in 2013–2014—an area equivalent to France, Germany, and the UK combined (WWF, 2015). However, plant-based foods and direct consumption contribute minimally to this, as more than 75% of soy is allocated to livestock feed rather than direct human or pet consumption (Fraanje and Garnett, 2020). In addition, meat protein production has been shown to have 6-17 times higher land requirement than soybean production (Reijnders and Soret, 2003). By contrast, chicken – while often viewed as a lower-impact protein – also relies heavily on soy and maize for fast growth, both of which compete with the human food system (Wilkinson, 2011).

### Greenhouse Gas Emissions

Beef and lamb-based diets led to dramatically higher CO_2_-equivalent emissions than poultry, plant-based, or veterinary formulations. These results mirror broader findings that 80% of agricultural GHG emissions arise from soil changes linked to deforestation, fertiliser use, and methane from ruminants (Pedrinelli *et al*., 2022). Although plant-based ingredients such as soy have drawn criticism for land use, their total emissions remain substantially lower per unit of protein. Knight (2023) emphasises that switching dogs to vegan diets could reduce total global GHG outputs by 5.8%.

### Acidifying and Eutrophying Emissions

Environmental degradation via nutrient pollution is another critical consequence of animal agriculture. Phosphates, commonly used as a key ingredient in pet food formulations to promote bone and dental health, are often sourced from phosphate rock during the manufacturing process which can lead to increases of excess phosphates into nearby water systems (Reijnders and Soret, 2003). They are also present in fertilisers, which are applied liberally to increase crop yield.

This study found plant-based diets yielded 7–16 times lower phosphate- and sulphur-based emissions than beef or lamb diets. These emissions contribute to acid rain and aquatic eutrophication, with long-term impacts on biodiversity (Chislock *et al.,* 2013). During the agricultural process, phosphate ions (PO_4_^3-^) are released into the environment, depleting soil richness and causing an overabundance of algae and plant growth (Scarborough *et al*., 2023), leading to the decomposition of plants, release of CO_2_, and subsequent water pH reduction and ocean acidification (National Oceanic and Atmospheric Association, 2024). Food production is the single largest contributor to global ocean and freshwater eutrophication, around 78% and 95%, respectively, of which most is captured at the farm stage (Poore and Nemecek, 2018). Similarly, Anthropocene activity contributes significantly to acidification of soil and water sources. This interferes with terrestrial and aquatic ecosystems by damaging plant life, degrading soil quality, and acidifying large bodies of water (European Environmental Agency, 2020). Jarosch, Bach and Finkbeiner (2024) previously reported that wet vegan foods still outperformed meat-based diets in eutrophication, though dry diets remain more sustainable overall.

### Freshwater Withdrawal

Water stress continues to increase in countries around the world, with 70% or more of global freshwater being used for agriculture (Knight, 2023; Tony Fujs and Haruna Kawshiwase, 2023). Beef and lamb required the most water per 1000kcal, with plant-based diets using the least. (Mekonnen and Hoekstra, 2011) previously found that water for crop production dwarfs all other uses, with wheat and maize among the most water-intensive crops. The global water footprint of crop production in the period 1996–2005 was 7404 Gm^3^ yr^−1^, with wheat making up 15% of this. Other crops with a large total water footprint are rice (992 Gm^3^ yr^−1^) and maize (770 Gm^3^ yr^−1^). However, whilst livestock drink little water directly, feed crop irrigation accounts for most of the sector’s water use (Pimentel and Pimentel, 2003). They estimate the production of forage and grain was estimated to use 100,000L of water to produce 1kg of fresh beef.

### But what about by-products?

Traditionally, dogs were fed table scraps, which had a lower impact on the environment (Su, Martens and Enders-Slegers, 2018). However, the rise in popularity of “premium” dog foods, which increasingly incorporates parts of the animal fit for human consumption, has created feed-food competition (Okin, 2017). Many of the meat-based protein sources analysed in this study were derived from meat meals rather than fresh meat, most of which are classified as animal by-products (ABPs), likely sourced as offcuts from the human food chain. These by-products are often considered a more sustainable option than prime meat cuts (Pingali *et al*., 2023), as they help reduce waste and make use of parts that might otherwise be discarded (Alexander *et al*., 2020). However, recent evidence suggests that the environmental benefits of by-products may be overstated. Knight (2023) reported that by-products not fit for human consumption require 1.35 times more livestock animals per unit of usable protein, compared to lean meat. Rushforth and Moreau (2013) similarly found that using offal in pet food may be less efficient than using muscle meat, due to lower digestibility and nutrient density. One advantage of using animal by-products is their potential to divert waste from landfill and improve protein recovery across the food system. But whilst ABP-based diets typically generate lower greenhouse gas emissions than whole-meat formulations, their emissions remain higher than those of plant-based diets. In our analysis, meat and bone meal—a common by-product—was among the highest-impact ingredients, with land use values of 327.70m², and CO₂-equivalent emissions of 99.78kg per 1000 kcal as methane and nitrous oxide from livestock, along with emissions from rendering processes, continue to contribute to their footprint. Likewise, although water use for rendering is relatively low, most of the water demand arises upstream from feed crop cultivation and animal husbandry, placing ABP diets above plant-based ones in terms of freshwater use. Moreover, some by-products used in pet food could otherwise be repurposed for human or industrial applications (Martens, Su and Deblomme, 2019), further complicating sustainability claims. The concept of using “waste” ingredients in pet food must therefore be critically assessed against the full lifecycle and opportunity cost of those inputs in other industries.

### Is a circular food system the answer?

Globally, 45% of the world’s habitable land is dedicated to agriculture, and of that, 70–85% is used to support livestock production (Van Zanten *et al*., 2018; Knight, 2023). This includes not only grazing grasslands, but considerable areas of cropland used to grow livestock feed. With a growing global population comes an increasing food demand, and the concept of a more circular food system is gaining traction (Van Selm *et al*., 2022), as well as emphasising the complementary roles of plants and livestock in sustainable food production (Godfray *et al*., 2010; Van Kernebeek *et al*., 2016; Van Zanten, Van Ittersum and De Boer, 2019). Van Zanten *et al*. (2018) found that compared to excluding ABPs, a system where livestock consume biomass unsuited for direct human consumption could free up to one quarter of global arable land. This could also provide around 9–23g of daily protein needs, per capita, although animal consumption above this level would require human-edible crops to be redirected to livestock. Schader *et al*. (2015) supported this and demonstrated that feeding animals primarily on grasslands and food production by-products, rather than human-edible concentrates, could decrease land use by 26% and cut GHG emissions by 18%.

While trade-offs remain, integrating these principles into pet food production aligns with broader efforts to enhance sustainability without increasing competition with human food supplies. In one study, using and marketing a pet food as “upcycled” enhanced its image, especially in budget-friendly markets (Ye *et al*., 2022). However, even though pet owners saw upcycled food as sustainable, this didn’t influence their buying behaviour as much as the perceived quality of the food. In addition to other available studies reviewing the topic, despite public hesitancy towards moving to plant-based dog food, the current authors have previously shown that they can indeed be as nutritionally complete as meat-based foods (Brociek *et al*., 2024).

### Study limitations

The impact values used in this study are at the end at the point of retail (Pedrinelli *et al*., 2022) and so only account for the production, manufacturing and distribution of the food. Therefore, additional environmental impacts of waste (food waste, animal waste and packaging end of life), previously explored (Swanson *et al*., 2013; Yavor, Lehmann and Finkbeiner, 2020), have not been accounted for. Modelling the environmental impacts of ingredients is limited by data availability, requiring assumptions for certain components. In some cases, our database lacked specific impact values, necessitating estimations based on analogous ingredients, which may not precisely represent current production methods. Additionally, as noted by Jarosch et al. (2024), assumed values may not scale linearly with global warming potential, introducing further uncertainty. While extreme miscalculations are unlikely, these limitations highlight the need for improved ingredient-specific data to enhance model reliability.

## Conclusion

Our findings re-confirm those of studies in other countries that plant-based dog foods are significantly more environmentally sustainable than traditional meat-based formulations. Though poultry- and veterinary-based foods showed lower impact than those produced with beef or lamb as the main constituents, they remain substantially less sustainable than plant-based alternatives. Use of animal by-products in such estimations, whilst helping to reduce food waste, do not get close to reducing the gap in environmental sustainability between meat- and plant-based foods. With rising global food demand and growing pet ownership, shifting toward lower-impact pet food ingredients will be essential in reducing the sector’s ecological paw print.

## Author contributions

R.A.B designed the research; R.A.B conducted the research; R.A.B and D.S.G analysed the data; D.S.G gained the funding and supervised the project; R.A.B and D.S.G co-wrote the manuscript. Both authors critically evaluated the paper and take responsibility for its final content.

## Acknowledgements and Financial Disclosure

The authors would like to thank Dr Jack Bobo, former Director of The Food Systems Institute, University of Nottingham (<u>Food Systems Institute</u>), for his advice whilst preparing the final version of the manuscript. This study was funded by the Biotechnology and Biological Sciences Research Council (BBSRC) as part of the University of Nottingham DTP PhD studentship awarded to R.A.B (Grant code: RS86P5).

